# The Enigma of UV Stress Granules: Research Challenges and New Perspectives

**DOI:** 10.1101/2022.10.11.511743

**Authors:** Alexandra J. Cabral, Danielle C. Costello, Natalie G. Farny

**Affiliations:** Department of Biology and Biotechnology, Worcester Polytechnic Institute, Worcester, MA, USA

**Author notes:** **Correspondence:** Natalie G. Farny, PhD.

**Keywords:** Stress granules, ultraviolet radiation (UV), biomolecular condensation, poly(A)+ RNA, cell cycle, neurodegeneration

## Abstract

Stress granules (SGs) are non-membrane bound cytoplasmic condensates that form in response to a variety of different stressors. Canonical SGs are thought to have a cytoprotective role, reallocating cellular resources during stress by activation of the integrated stress response (ISR) to inhibit translation and avoid apoptosis. However, different stresses result in compositionally distinct, non-canonical SG formation that is likely pro-apoptotic, though the exact function(s) of both SGs subtypes remain unclear. A unique non-canonical SG subtype is triggered upon exposure to ultraviolet (UV) radiation. While it is generally agreed that UV SGs are *bona fide* SGs due to their dependence upon the core SG nucleating protein Ras GTPase-activating protein-binding protein 1 (G3BP1), the localization of other key components of UV SGs are unknown or under debate. Further, the dynamics of UV SGs are not known, though unique properties such as cell cycle dependence have been observed. This Perspective compiles the available information on SG subtypes and on UV SGs in particular in an attempt to understand the formation, dynamics, and function of these mysterious stress-specific complexes. We identify key gaps in knowledge related to UV SGs, and examine the unique aspects of their formation. We propose that more thorough knowledge of the distinct properties of UV SGs will lead to new avenues of understanding of the function of SGs, as well as their roles in disease.

## 1 Introduction

Stress granules (SGs) are evolutionarily conserved cytoplasmic biomolecular condensates that form in response to a variety of environmental stressors (Nover et al., 1989; Kedersha et al., 1999, 2002) Since the discovery of SGs in mammalian cells in 1999 (Kedersha et al., 1999)an ever-broadening array of stressors have been identified as inducers of SGs. While it is known that different stressors can give rise to compositionally and functionally distinct SGs (Aulas et al., 2017; Advani and Ivanov, 2020), our understanding of the components of SG subtypes, and their functional consequences, is understudied. The study of SGs is of intense interest due to the role SGs likely play in various diseases, especially protein aggregation diseases such as Alzheimer’s Disease (AD) and Amyotrophic Lateral Sclerosis (ALS) (Liu-Yesucevitz et al., 2010; Li et al., 2013; Ash et al., 2014; Boyd et al., 2014; Wolozin, 2014; Wolozin and Ivanov, 2019) Because SG function may be determined both by composition and by the environmental conditions under which they form, gaining insights into compositionally distinct SG subtypes will be important for understanding and manipulating SG formation in the context of disease.

One of the most distinct and least understood SG subtypes are those induced by ultraviolet radiation (UV). In countless reports and reviews of SG biology, UV is listed among the litany of stresses that induce SG formation. Humans experience UV daily in the environment, and its genotoxic and carcinogenic effects have been well studied (Roy, 2017). However, published evidence documenting the composition, dynamics, and function of UV-induced SGs is surprisingly limited. We do not yet understand the mechanism(s) driving UV SG assembly, nor the functional consequences of their formation. In this Perspective, we assemble the available evidence on the biology of UV SGs. We believe that this unique SG subtype, which unlike most SGs seems to lack mRNA, is also the only known example of G1-specific cell cycle-controlled SG formation. We propose that the unique properties of UV SGs represent an untapped source for new hypotheses to contribute broadly to the fields of SG biology, aggregation-mediated disease, and biomolecular condensation.

## 2 Understanding SG Subtypes

### 2.1 Canonical SGs

As one of the earliest discovered and most broadly studied SG types, SGs induced by acute high-dose exposure to sodium arsenite have become the canon by which SGs are defined (Kedersha et al., 1999, 2002; Aulas and vande Velde, 2015; Aulas et al., 2017; Advani and Ivanov, 2020; Riggs et al., 2020). So-called “canonical” or arsenite-like SGs – caused by many stresses including heat shock (Kedersha et al., 1999), thapsigargin (Aulas et al., 2017), and bisphenol A (Fay et al., 2021) among others – are associated with the inhibition of global protein synthesis and the preferential translation of stress-induced transcripts(Harding et al., 2000a). Bulk translation is inhibited through two main pathways(Advani and Ivanov, 2019): 1) the phosphorylation of the alpha subunit of eukaryotic initiation factor 2 alpha (P-eIF2α), which occurs in mammalian cells via one of four serine/threonine kinases: (heme-regulated eIF2α kinase (HRI) (McEwen et al., 2005), protein kinase R (PKR) (Srivastava et al., 1998), PKR-like ER kinase (PERK) (Harding et al., 2000b) and general control nonderepressible 2 (GCN2)(Wek et al., 1995); and 2) the inhibition of the mammalian target of rapamycin (mTOR)(Proud, 2019). The former mechanism inhibits recycling of the eIF2/tRNA_i_^Met^/GTP ternary complex required for translation initiation (Wek, 2018), while the latter mechanism results in hypophosphorylated eIF4E binding protein (4E-BP) which binds eIF4E and inhibits translation (Proud, 2019). When translation is inhibited, 48S translation pre-initiation complexes and untranslated mRNAs accumulate and aggregate in the cytoplasm, favoring a liquid-liquid phase separation (LLPS) event (Guillén-Boixet et al., 2020; Sanders et al., 2020; Yang et al., 2020) and assembling into SGs. Canonical SGs are in dynamic equilibrium with polysomes, as drugs that inhibit polysome disassembly (e.g., emetine, cycloheximide) inhibit SG formation, while drugs that induce premature polysome disassembly (e.g., puromycin) promote SG assembly (Kedersha et al., 2000, 2002).

The first protein markers associated with SGs were the RNA-binding proteins TIA-1 and TIAR, as well as the cytoplasmic poly(A)-binding protein (PABPC1)(Kedersha et al., 1999). Soon thereafter, the Ras GTPase-activating protein-binding protein 1 (G3BP1) was identified as a key modulator of SG assembly (Tourrière et al., 2003). It is now firmly established that G3BP1 is a master regulator of SG assembly, and most known SG subtypes require G3BP1 or its close homolog G3BP2 for their assembly (Kedersha et al., 2016; Guillén-Boixet et al., 2020; Sanders et al., 2020; Yang et al., 2020). Related to their relationship with eIF2α-mediated translational control, canonical SGs contain poly(A)+ mRNAs as well as stalled translation 48S-preinitiation complexes including the eIF3 complex, eIF4G, and the small ribosomal subunit (Kedersha et al., 2002; Aulas et al., 2017) (Table 1).

**Table 1:**
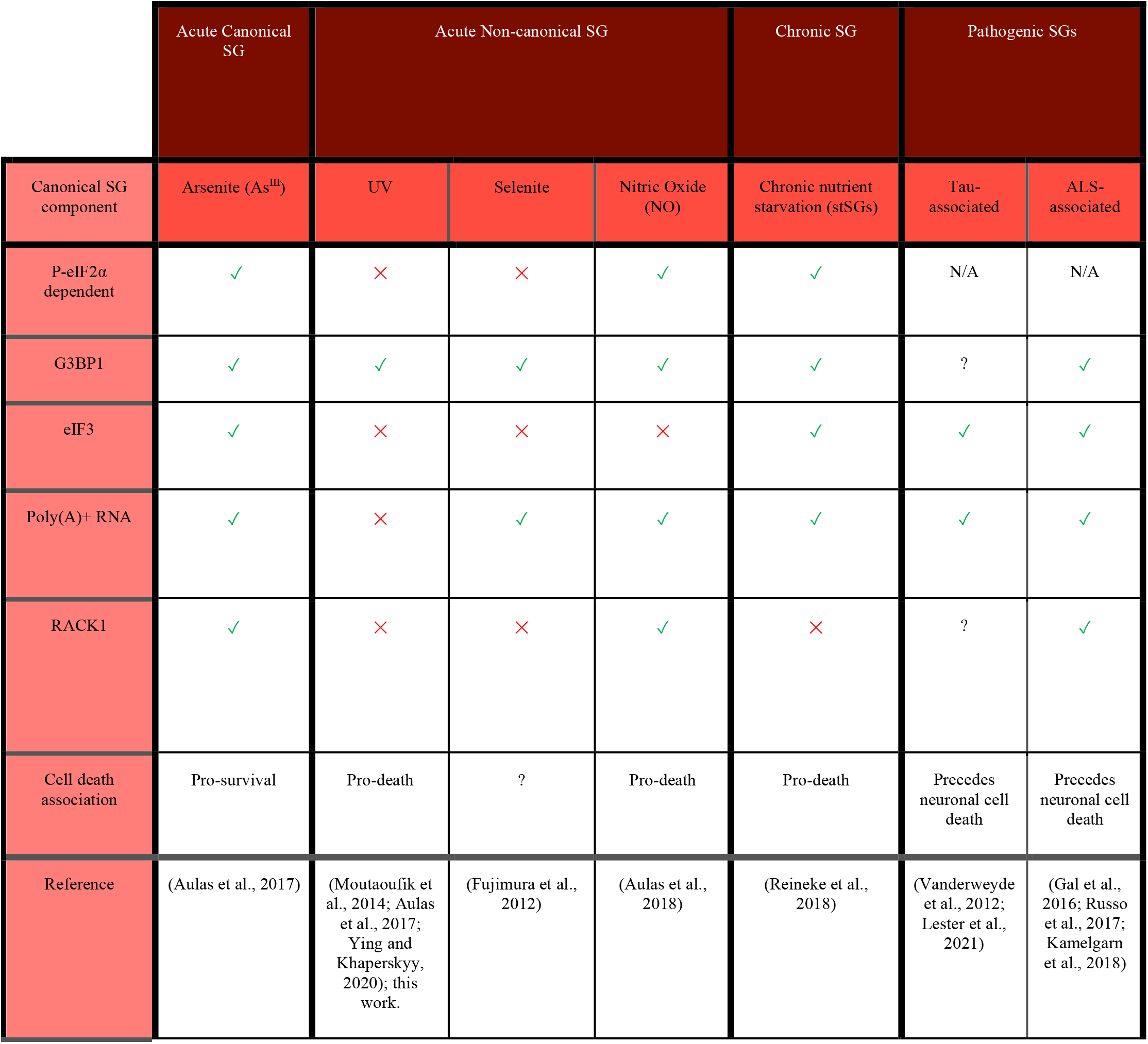
A comparison of key features of canonical, non-canonical, chronic, and pathogenic SG subtypes.

While the composition of canonical SGs have been intensively catalogued through affinity labeling and biochemical purification studies (Jain et al., 2016; Markmiller et al., 2018; Youn et al., 2019; Marmor-Kollet et al., 2020; An et al., 2022), the functional consequences of SG formation remain unclear. One proposed function for SGs is the modulation of apoptosis^12,13^(Arimoto et al., 2008; Arimoto-Matsuzaki et al., 2016; Park et al., 2020). In a now classic report, Arimoto et al. showed that SGs caused by overexpression of G3BP1 conferred resistance to GADD45-mediated apoptosis (Arimoto et al., 2008). They further demonstrated that hypoxia, which triggers SGs, protected cells from etoposide-mediated apoptosis. In both cases, they confirmed that sequestration of the signaling scaffold protein RACK1 at the SG was essential for protection from apoptosis (Arimoto et al., 2008). Other putative functions for SGs include mRNA triage (Kedersha et al., 2005) and stress-induced translational control (Dey et al., 2010); while SGs likely play a role, the idea that SG formation is absolutely essential or even largely responsible for such processes has largely been discredited (Mateju et al., 2020; Glauninger et al., 2022; Mateju and Chao, 2022).

SGs are dynamic. Canonical SGs rapidly exchange factors with the surrounding cytoplasm, though some factors exchange more rapidly than others (Kedersha et al., 2000). The extent to which factors move in and out of SGs is related to the structure of the granule, which appears to have a less dynamic core surrounded by a more dynamic shell (Wheeler et al., 2016). SGs that are more liquid-like also tend to be more dynamic, whereas those associated with gel or solid-like states (including some non-canonical and pathogenic subtypes, described below) are less dynamic (Kroschwald et al., 2015; Zhang et al., 2019).

### 2.2 Non-Canonical SGs

The so-called “non-canonical” SG is a category used to collect SGs that do not mirror the formation, composition or function of canonical SGs as primarily understood from studies of arsenite and heat shock SGs (Advani and Ivanov, 2020). This SG subtype was originally dubbed non-canonical based on the lack of the canonical SG protein eukaryotic initiation factor 3 (eIF3) (Advani and Ivanov, 2020; Hofmann et al., 2021) in SGs triggered by sodium selenite (Fujimura et al., 2012), which suggested that the 48S preinitiation complex was not recruited to these SGs. Like canonical SGs, non-canonical SGs form acutely (in 4 hours or less of stress), require G3BP1 or G3BP2 for their formation (Yang et al., 2020), and are associated with translational arrest, though they may or may not be associated with eIF2α phosphorylation depending upon the stress (Aulas et al., 2017; Advani and Ivanov, 2020; Hofmann et al., 2021). Non-canonical SGs are reported to form in response to many stresses including sodium selenite (Fujimura et al., 2012), nitric oxide (NO) (Aulas et al., 2018), the eIF4A inhibitor Rocaglamide A (RocA) (Aulas et al., 2017), hydrogen peroxide (Emara et al., 2012), and UV radiation (Kedersha et al., 1999; Pothof et al., 2009). There is broad variation within this non-canonical category as to which components do and do not localize at the SG, examples of which are described in Table 1. Others have attempted to further subdivide canonical and non-canonical SGs into designations including Types I, II and III (Hofmann et al., 2021), however there is no consensus and some conflicting characterizations among these classifications (Advani and Ivanov, 2020; Hofmann et al., 2021), and thus we will address all acute non-canonical SG subtypes holistically in this Perspective.

Non-canonical SGs are reported to be functionally distinct from canonical SGs in that they are believed to be cytotoxic and less dynamic (Advani and Ivanov, 2020). However, there are few published studies that experimentally validate these characterizations. Relative to cell death, the most thoroughly characterized non-canonical SGs are those caused by nitric oxide (NO). NO SGs were determined to be triggering a non-apoptotic and potentially necrotic cell death by assaying a combination of propidium iodide and trypan blue staining, ATP levels and Caspase-3 cleavage (Aulas et al., 2018). For other non-canonical stresses like selenite, RocA, and UV, a decrease in cell viability was associated with SGs, but a mechanism of death was not elucidated (Fujimura et al., 2012; Aulas et al., 2018). Interestingly, while canonical arsenite-induced SGs are believed to be anti-apoptotic (Arimoto et al., 2008), the inverse relationship does not appear to be a universal feature of non-canonical SG subtypes, in that the lack of sequestration of pro-apoptotic factors like RACK1 at non-canonical SGs cannot simply be interpreted to mean that those pro-apoptotic factors are therefore active and inducing apoptotic cell death associated with non-canonical SG formation. Relative to their dynamics, the only report of which we are aware that directly measured dynamic behavior of a protein component of a non-canonical SG is in the context of NO-induced SGs (Aulas et al., 2018). G3BP1 recovery to photobleached NO SGs was approximately 10% lower than in arsenite SGs (Aulas et al., 2018). To our knowledge, this summarizes the extent to which the cytotoxic and dynamic properties of acute non-canonical SGs have been directly assessed in the literature.

### 2.3 Chronic and Pathological SG Subtypes

The canonical and non-canonical labels are most frequently associated with acute SG formation. Chronic stresses and disease states are associated long-term SG formation. Chronic starvation induced SGs (stSGs) appear after prolonged (8-16 hours) starvation of glucose, serum, glutamine and pyruvate (Reineke et al., 2018). These stSGs require both G3BP1 and eIF2α phosphorylation, and are in dynamic equilibrium with polysomes. StGSs appear to be pro-apoptotic, as assessed by Annexin V staining (Reineke et al., 2018), but the dynamic properties of these stSGs remain unknown. Pathological SGs (pSGs) are chronic SGs that form in diseased cells and are hypothesized to seed the accumulation of irreversible and toxic disease aggregates. pSGs have been most closely associated to date with ALS and Alzheimer’s Disease (Vanderweyde et al., 2012, 2016; Ash et al., 2014; Aulas and vande Velde, 2015; Apicco et al., 2018; Zhang et al., 2019; Marmor-Kollet et al., 2020; An et al., 2022). Several excellent reviews have been written on the subject (Wolozin, 2014; Aulas and vande Velde, 2015; Wolozin and Ivanov, 2019), including one within this Research Topic (Rhine et al., 2022). Interestingly, stSGs and pSGs can contain many of the components of canonical granules including poly(A)+ RNA, eIF3, and eIF4G (Table 1), though unlike canonical SGs they are clearly associated with (Liu-Yesucevitz et al., 2010; Vanderweyde et al., 2012; Reineke et al., 2018) This observation raises the question of whether the categorization of SGs into canonical and non-canonical groups based primary on composition data will remain a useful proxy for the breadth of SG function as we continue to discover SGs in new contexts.

## 3 UV SGs are a unique non-canonical SG subtype

### 3.1 Discovery and early observations of UV SGs

UV SGs were first reported in 1999 by Kedersha et al. (as data not shown) as containing the SG marker proteins TIA-1, TIAR, and PABPC1 (Kedersha et al., 1999). The first published evidence of UV-induced cytoplasmic aggregation was in 2005, when Teixeira et al. were investigating processing bodies (PBs, a cytoplasmic mRNP complex associated with mRNA decay (Luo et al., 2018)), in the budding yeast *S. cerevisiae* (Teixeira et al., 2005). Then in 2008, UV-induced poly(A)+ RNA-containing bodies were reported in *S. cerevisiae* by Gaillard et al., but were not thought to be *bona fide* SGs due to the lack of co-localization of putative SG homologs (Gaillard and Aguilera, 2008). At that time however, SGs had not yet been discovered in *S. cerevisiae*, and thus core SG markers had not been established in that species.

Finally in 2009, Pothof et. al. investigated the DNA damage response (DDR) to UV in HeLa cells, and discovered that the cytoplasmic condensates they observed in response to UV are SGs, based on the co-localization of TIA-1 and Ago2, and that UV SGs may be involved in microRNA-mediated silencing in response to DNA damage (Pothof et al., 2009). Several more recent characterize UV SGs as non-canonical SGs lacking eIF3 and eIF4G (Table 1) (Moutaoufik et al., 2014; Aulas et al., 2017; Ying and Khaperskyy, 2020). We confirm here that UV SGs also lack the pro-apoptotic scaffolding protein RACK1, 4 hours after UV treatment of U2OS human osteosarcoma cells (Figure 1A). Like other non-canonical subtypes, UV SGs are G3BP-dependent, eIF2α-independent, and cytotoxic (Pothof et al., 2009; Moutaoufik et al., 2014; Aulas et al., 2017; Ying and Khaperskyy, 2020). Surprisingly however, there are conflicting reports about whether UV SGs contain poly(A)+ RNA (Moutaoufik et al., 2014; Aulas et al., 2017).

**Figure 1:**
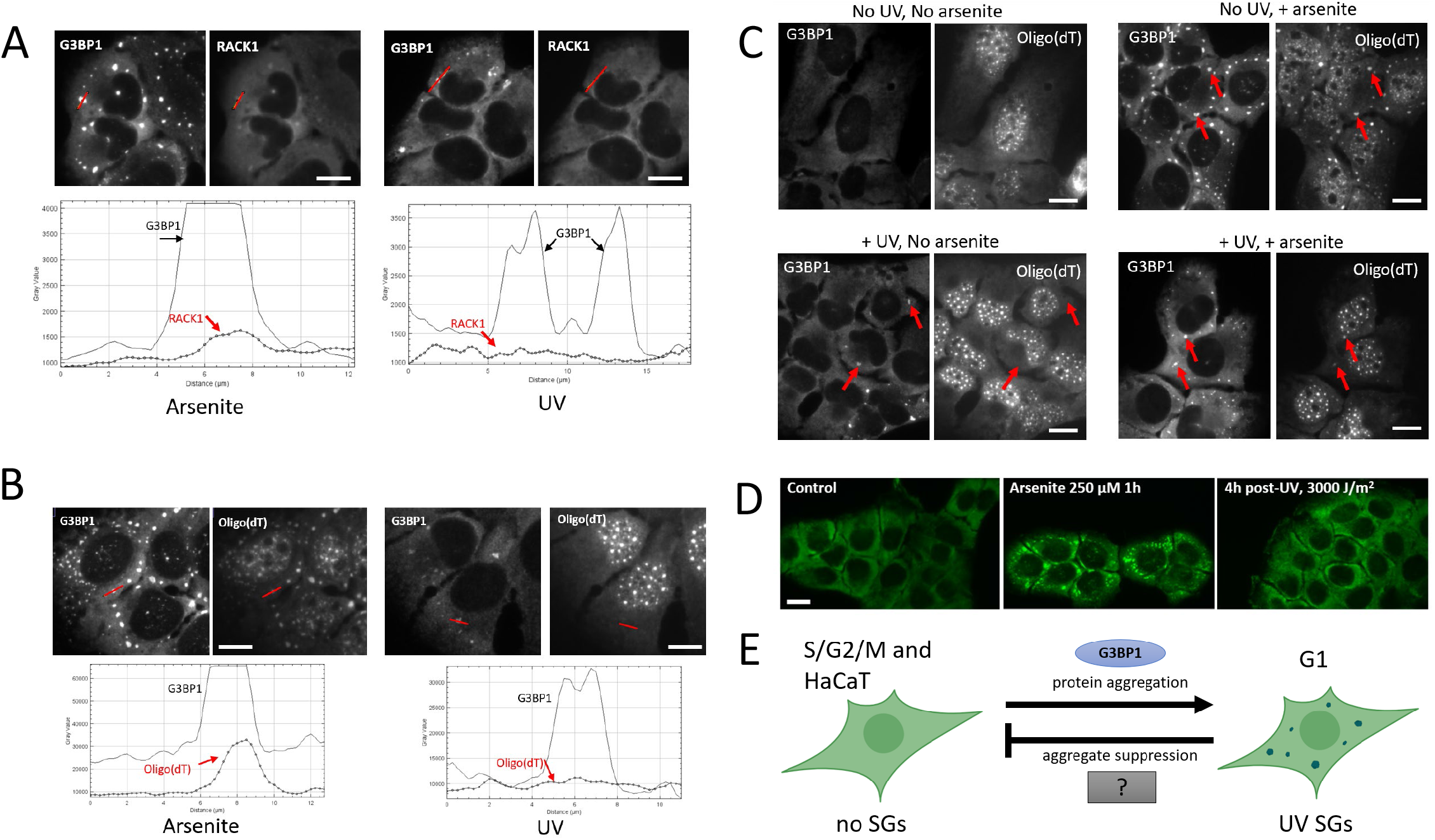
Composition and formation of UV SGs. A. RACK1 does not localize to UV-induced SGs. U2OS treated with arsenite (500 µM, left panels) or UV (15 J/m^2^, then assayed at 4 hours post-UV, right panels) and co-stained with antibodies to G3BP1 and RACK1. Profile intensity plots over the red line from each image were compiled using ImageJ. B. U2OS treated with arsenite (500 µM, left panels) or UV (15 J/m^2^, then assayed at 4 hours post-UV, right panels) and co-stained with antibodies to G3BP1 and Oligo(dT). Profile intensity plots over the red line from each image were compiled using ImageJ. C. Poly(A)+ RNA SGs can form after UV treatment. U2OS were untreated or treated with UV (15 J/m^2^) for 3 hours, then arsenite (500 µM) was added where indicated for a 1 hour, then cells were fixed and co-stained with antibodies to G3BP1 and Oligo(dT) FISH. Red arrows point to prominent SGs in each treatment condition. D. HaCaT cells do not form UV SGs. HaCaT cells were exposed to arsenite 250 µM for 1 hour, or exposed to 3000 J/m^2^ UV (4 hours post-exposure), then stained with antibody for G3BP1 to assess SG formation. Detailed methods for all panels are available as supplementary material online. E. A model for UV SG suppression. See Discussion section for details.

### 3.2 Are UV SGs RNA Granules?

It was widely assumed in the early years of their discovery that all SGs contained mRNA. SGs were defined as mRNA-containing bodies or mRNP aggregates in many articles and reviews (Anderson and Kedersha, 2006; Riggs et al., 2020; Hofmann et al., 2021). In 2014, Moutaoufik et al. observed the colocalization of poly(A)+ RNA at UV SGs by co-localizing an oligo(dT) probe with the Fragile X Mental Retardation Protein (FMRP) as a SG marker in NIH-3T3 cells 18 hours post-UV exposure (Moutaoufik et al., 2014). This report is, to our knowledge, the first published account of RNA localization to UV SGs in a mammalian cell type. However, a study published in 2017 by Aulas et al. found no poly(A)+ RNA localization at UV SGs, using G3BP1 as a SG marker in Hap1 human haploid cells, two hours post-UV (Aulas et al., 2017)

To independently validate the observation of poly(A)+ RNA at UV SGs, we treated U2OS cells with 15 J/m^2^ UVC, and used fluorescence in situ hybridization (FISH) to examine poly(A)+ RNA localization to UV SGs using G3BP1 as a SG marker. We find, as did Aulas et al. (2017), that poly(A)+ RNA is not enriched at UV SGs at four hours post-UV exposure. (Figure 1B). It is tempting to speculate that perhaps UV light degrades cellular pools of mRNA, which could explain why poly(A)+ RNA FISH signal does not accumulate at UV SGs. However, the literature actually suggests that UV stabilizes and inhibits the degradation of mRNAs (Burgess et al., 2011). mRNAs are sequestered in the nucleus (Burgess et al., 2011) which may contribute to the loss of signal at cytoplasmic UV SGs. PABP1 and PABP4 also relocalize to the nucleus, however the poly(A)+ RNA nuclear retention and PABP localization appear to be independent, which further supports a dissociation of PABPs from their RNA targets (Burgess et al., 2011). Although it is reported UV can stabilize mRNAs, it is also known that UV induces reactive oxygen species and that oxidized RNAs can be targeted for degradation^43^, meaning that some mRNA degradation may also be possible.

To determine whether it was likely that mass degradation of mRNA was responsible for the lack of poly(A)+ RNA we observed at UV SGs, we treated U2OS with UV, then used arsenite to induce SGs at three hours post-UV exposure. We observe that arsenite SGs induced in UV irradiated cells contain poly(A)+ RNA, though the intensity of localization is decreased compared to arsenite SGs in non-UV treated cells (Figure 1C). Therefore, we submit that mass degradation of mRNA cannot fully explain the lack of poly(A)+ RNA signal at UV SGs, as we show it is possible to assemble poly(A)+-containing SGs after UV treatment. We conclude that poly(A)+ RNAs are specifically excluded from UV SGs, though the mechanism of this exclusion is unknown. The results also do not preclude the possibility that deadenylated mRNAs, non-coding mRNAs and/or small RNAs may still be present, and this possibility warrants further investigation. If, however, it is the case that UV SGs are not RNA granules, it would mean that UV SGs do not play any role in the RNA-associated putative functions of SGs such as mRNA triage and translational control.

The finding that UV SGs are depleted of mRNA may also have implications for the assembly of SGs in general. Numerous reports have documented the role of RNA in driving LLPS during SG formation (van Treeck et al., 2018; Guillén-Boixet et al., 2020; Yang et al., 2020; Matheny et al., 2021). If indeed UV SGs do not contain mRNA, then one concludes that the interactions that drive UV SG assembly, unlike most other SG subtypes, must be primarily protein-driven. The lack of facilitation by RNA in LLPS of UV SG formation may also explain why UV SGs have been reported to be smaller and less numerous than other SG subtypes (Moutaoufik et al., 2014). Another reasonable prediction is that UV SGs may be more solid or gel-like than canonical or mRNA-containing SGs, as RNA is known to increase the liquid-like properties of biomolecular condensates (Guillén-Boixet et al., 2020; Yang et al., 2020; Bevilacqua et al., 2022). If indeed UV SG assembly is primarily protein-driven, then UV SGs may be a useful model for the specific study of essential protein-protein interactions within SGs. The dynamic properties of UV SGs have not yet been measured, but will no doubt be the topic of future investigation.

### 3.3 UV SG Suppression

Because SGs have been implicated in seeding pathological protein aggregation (Wolozin and Ivanov, 2019), mechanisms of SG suppression are of intense interest. For example, SG induced by arsenite, a strong oxidizer, can be suppressed by strong antioxidants such as N-acetylcysteine (Szaflarski et al., 2016; Aulas et al., 2018) as well as by preconditioning that likely enhances cellular resilience to oxidative stress (Glass and Wente, 2019; Fay et al., 2021). SGs are also known to be suppressed in mitotic cells, because stalled translation elongation prohibits polysome disassembly in M phase (Gilad et al., 2007). While assessing the DNA damage response, Pothof et al. noted that UV-induced SGs did not form in cells that were positive for cyclin A, a marker of S phase (Pothof et al., 2009). By synchronizing their cells prior to irradiation, it was determined that UV SGs only appeared in cells that were irradiated in G2, passed through mitosis, and were in G1 phase at the assay timepoint. This observation of cell cycle dependence likely explains why only ∼10-30% of cells in an asynchronous culture form UV SGs (Pothof et al., 2009; Aulas et al., 2017). The mechanism for the G1 specificity of UV SGs is unknown, however the ATM and ATR DNA damage checkpoint kinases were not required for UV SG formation (Pothof et al., 2009).

The observation of cell cycle dependence of UV SGs was confirmed in 2014 by Moutaoufik et. al., who discovered that a dose of 10 J/m^2^ UV resulted in complete G_1_ arrest (Moutaoufik et al., 2014).These researchers used FACS profiling to assess the cell cycle, while concurrently assessing SG formation by immunofluorescence microscopy, and reported that G1 arrest correlated with the highest SG formation, and resumption of the cell cycle corresponded with SG clearance. Although this implies UV SGs may be dissolvable, it is also possible that the cells with UV SGs underwent cell death, leaving behind SG-free cells to resume the cell cycle. Thus, suppression of UV SGs in S and G2 phases is a unique feature of UV SGs.

Because skin is the organ most directly affected by UV, we decided to examine UV SG formation in an untransformed human keratinocyte cell line, HaCaT. To our great surprise, we were completely unable to induce UV SGs in HaCaT cells. We applied increasing amounts of UV, from our starting dose of 15 J/M^2^ up to 3000 J/M^2^, and observed zero UV SG formation in HaCaTs by G3BP1 staining (Figure 1D). We do not yet know if this intriguing observation of UV SG suppression is a common property of all keratinocytes or specific to HaCaT, nor do we know whether it is related to the mechanisms of G2, S, and M phase suppression described above. We should note here that all studies of UV SGs reported to date, including the data reported herein, use UVC (254 nm) light sources. UVC is well known to cause damage to human tissues (Roy, 2017), but does not penetrate the Earth’s atmosphere. It remains unknown whether UVA (400-315 nm) and UVB (315-280 nm) rays can trigger UV SG formation in any cell type, including keratinocytes. However, UVA causes significantly more reactive oxygen species in human tissues than other UV wavelengths (Karran and Brem, 2016), which would cause an increase in cellular stress over other wavelengths.

## 4 Discussion

The observation that UV SGs are RNA-depleted, protein-driven, and cell cycle dependent bodies makes them unique among known SG subtypes, and therefore they are fascinating research targets. Because UV SGs do not appear to contain much if any RNA (Figure 1B), we hypothesize that they primarily represent the effects of protein-protein interaction-driven condensation. In this way, UV SGs are a platform for studying the protein-specific drivers and suppressors of SG phase separation. The observations of SG suppression in S/G2/M phases, as well as in keratinocytes (Figure 1D), coupled with the apparent lack of poly(A)+ RNA (Figure 1B), informs our model for UV SG suppression (Figure 1E). In response to UV, which may cause nuclear retention and/or RNA-binding protein dissociation from RNA (Burgess et al., 2011), the protein drivers of SG formation including G3BP1/2 are just barely capable of surpassing a critical threshold and achieving protein-driven phase separation. In S/G2/M phases, and in certain UV-prone cell types such as keratinocytes, yet unknown cytoplasmic factors suppress phase separation by shifting the equilibrium to disfavor protein-driven condensation. This model is supported by the observation that G3BP1 condensation is significantly facilitated by the presence of RNA both *in vitro* and *in vivo* (Guillén-Boixet et al., 2020; Yang et al., 2020). Thus, in the absence of RNA, cytoplasmic factors may play a greater role in favoring or disfavoring biomolecular condensation in a way that would not be detectable if RNA were there to push the balance toward condensate formation. The “mystery SG suppressor” may not be a single gene, but a set of cytoplasmic environmental conditions associated with cell cycle progression. Cross-referencing proteomic, transcriptomic, and metabolomic differences between G1 and the rest of the cell cycle with similar data from keratinocytes is a starting point to identify the hidden factor(s) regulating UV SG suppression.

The unique nature of UV SGs may have revealed to us a path forward to identify suppressors of other protein aggregates. Many reviews have built powerful arguments for the relationship of SGs to protein aggregation diseases including ALS and AD (Li et al., 2013; Ash et al., 2014; Wolozin, 2014; Aulas and vande Velde, 2015; Wolozin and Ivanov, 2019). We suggest that the identification of UV SG suppressors, as modulators of a protein-driven condensation process, represent a new frontier for research in neurodegeneration. If SGs can become pathological and can seed toxic aggregation of proteins like tau and TDP-34, then suppressors of SG formation may prevent or even reverse these seeds and thereby prevent disease aggregates from forming. It is also worth noting that the affected cells in neurodegenerative disease – neurons – are perpetually frozen in G0; UV SGs are repressed in S/G2/M phases. If we can understand the cellular conditions underlying cell cycle-dependent UV SG suppression, we could potentially apply this knowledge to create a G2-like state in neurons and thereby suppress pathological aggregates. In light of their potential to illuminate new paths for protein aggregation disease research, addressing these questions about an oft-neglected SG subtype takes on a fresh urgency.

Many key gaps in knowledge about non-canonical SGs in general, and UV SGs in particular, remain to be addressed. It is still entirely unclear how UV irradiation triggers SG formation in the first place. The lack of poly(A)+ RNA suggests that perhaps G3BP1, and other RNA-binding proteins like PABPs (Burgess et al., 2011) may dissociate from their mRNA targets in response to UV stress. But the pathways that would lead to such dissociation are unknown. Identifying the components of UV SGs is a key step to understanding these unusual SGs. Biochemical purification strategies and proximity labeling studies have enabled us to understand the proteomic and transcriptomic landscapes of canonical SGs (Jain et al., 2016; Khong et al., 2017; Markmiller et al., 2018; Namkoong et al., 2018; Padrón et al., 2019; Youn et al., 2019; Marmor-Kollet et al., 2020; Matheny et al., 2021; Vu et al., 2021; An et al., 2022). However, parallel studies on UV SGs, or any non-canonical subtype, remain to be performed. The dynamics of UV SGs have not been directly measured, and while the evidence presented here leads us to predict that UV SGs have a more solid or gel-like than liquid like behavior, for now their dynamic nature is unclear. It further has not been investigated whether UV SGs, like canonical subtypes, are also organized into core and shell subdomains, or whether their substructure is substantially distinct form known SGs. Finally, while we haven’t addressed the topic here, most canonical and non-canonical SGs are believed to be reversible and to dissolve upon the resolution of stress, facilitated by the autophagy pathway (Buchan et al., 2013). The mechanisms of UV SG dissolution, if they are in fact reversible, remain unknown.

## 7 Conflict of Interest

The authors declare that the research was conducted in the absence of any commercial or financial relationships that could be construed as a potential conflict of interest.

## 8 Author Contributions

N.G.F. conceived of the project. All authors participated in experimentation that led to the results reported in Figure 1. A.J.C. assembled Table 1. N.G.F. assembled Figure 1. A.J.C and D.C.C wrote the initial draft. N.G.F. edited the draft. All authors participated in the final preparation of the manuscript for publication. N.G.F. provided supervisory and financial support.

## 9 Funding

This research is supported by NIH R03AG077140 to N.G.F., and new faculty start up funds from WPI to N.G.F.

## 10 Acknowledgments

The authors thank Nancy Kedersha, Paul Anderson, and Pavel Ivanov (Brigham and Women’s Hospital, Boston, MA) for U2OS cells; Louis Roberts (WPI) for HaCaT cells; members of the Farny lab for helpful discussions.

## 11 Supplementary Material

A supplementary file containing Materials and Methods and Supplementary References accompanies this manuscript

## Notes

### Competing Interest Statement

The authors have declared no competing interest.

## References

Advani VM, Ivanov P (2019) Translational Control under Stress: Reshaping the Translatome. BioEssays 41:1900009

Advani VM, Ivanov P (2020) Stress granule subtypes: an emerging link to neurodegeneration. Cellular and Molecular Life Sciences 77:4827–4845

An H, Litscher G, Watanabe N, Wei W, Hashimoto T, Iwatsubo T, Buchman VL, Shelkovnikova TA (2022) ALS-linked cytoplasmic FUS assemblies are compositionally different from physiological stress granules and sequester hnRNPA3, a novel modifier of FUS toxicity. Neurobiol Dis 162:105585

Anderson P, Kedersha N (2006) RNA granules. J Cell Biol 172:803–808.

Apicco DJ, Ash PEA, Maziuk B, LeBlang C, Medalla M, al Abdullatif A, Ferragud A, Botelho E, Ballance HI, Dhawan U (2018) Reducing the RNA binding protein TIA1 protects against tau-mediated neurodegeneration in vivo. Nat Neurosci 21:72.

Arimoto K, Fukuda H, Imajoh-Ohmi S, Saito H, Takekawa M (2008) Formation of stress granules inhibits apoptosis by suppressing stress-responsive MAPK pathways. Nat Cell Biol 10:1324.

Arimoto-Matsuzaki K, Saito H, Takekawa M (2016) TIA1 oxidation inhibits stress granule assembly and sensitizes cells to stress-induced apoptosis. Nat Commun 7:10252 Available at:

Ash PEA, Vanderweyde TE, Youmans KL, Apicco DJ, Wolozin B (2014) Pathological stress granules in Alzheimer’s disease. Brain Res 1584:52–58.

Aulas A, Fay MM, Lyons SM, Achorn CA, Kedersha N, Anderson P, Ivanov P (2017) Stress-specific differences in assembly and composition of stress granules and related foci. J Cell Sci 130:927– 937.

Aulas A, Lyons SM, Fay MM, Anderson P, Ivanov P (2018) Nitric oxide triggers the assembly of “type II” stress granules linked to decreased cell viability. Cell Death Dis 9:1129 Available at:

Aulas A, vande Velde C (2015) Alterations in stress granule dynamics driven by TDP-43 and FUS: a link to pathological inclusions in ALS?. Frontiers in Cellular Neuroscience 9:423

Bevilacqua PC, Williams AM, Chou H-L, Assmann SM (2022) RNA multimerization as an organizing force for liquid–liquid phase separation. RNA 28:16–26

Boyd JD, Lee-Armandt JP, Feiler MS, Zaarur N, Liu M, Kraemer B, Concannon JB, Ebata A, Wolozin B, Glicksman MA (2014) A high-content screen identifies novel compounds that inhibit stress-induced TDP-43 cellular aggregation and associated cytotoxicity. J Biomol Screen 19:44–56.

Buchan JR, Kolaitis R-M, Taylor JP, Parker R (2013) Eukaryotic stress granules are cleared by autophagy and Cdc48/VCP function. Cell 153:1461–1474.

Burgess HM, Richardson WA, Anderson RC, Salaun C, Graham S v, Gray NK (2011) Nuclear relocalisation of cytoplasmic poly (A)-binding proteins PABP1 and PABP4 in response to UV irradiation reveals mRNA-dependent export of metazoan PABPs. J Cell Sci 124:3344–3355.

Dey S, Baird TD, Zhou D, Palam LR, Spandau DF, Wek RC (2010) Both transcriptional regulation and translational control of ATF4 are central to the integrated stress response. J Biol Chem 285:33165–33174.

Emara MM, Fujimura K, Sciaranghella D, Ivanova V, Ivanov P, Anderson P (2012) Hydrogen peroxide induces stress granule formation independent of eIF2α phosphorylation. Biochem Biophys Res Commun 423:763–769.

Fay MM, Columbo D, Cotter C, Friend C, Henry S, Hoppe M, Karabelas P, Lamy C, Lawell M, Monteith S, Noyes C, Salerno P, Wu J, Zhang HM, Anderson PJ, Kedersha N, Ivanov P, Farny NG (2021) Bisphenol A promotes stress granule assembly and modulates the integrated stress response. Biol Open 10:bio057539

Fujimura K, Sasaki AT, Anderson P (2012) Selenite targets eIF4E-binding protein-1 to inhibit translation initiation and induce the assembly of non-canonical stress granules. Nucleic Acids Res 40:8099–8110.

Gaillard H, Aguilera A (2008) A novel class of mRNA-containing cytoplasmic granules are produced in response to UV-irradiation. Mol Biol Cell 19:4980–4992.

Gal J, Kuang L, Barnett KR, Zhu BZ, Shissler SC, Korotkov K v, Hayward LJ, Kasarskis EJ, Zhu H (2016) ALS mutant SOD1 interacts with G3BP1 and affects stress granule dynamics. Acta Neuropathol 132:563–576.

Gilad S, Nancy K, Orna E-S (2007) Ribosomal Slowdown Mediates Translational Arrest during Cellular Division. Mol Cell Biol 27:6639–6646 Available at:

Glass L, Wente SR (2019) Gle1 mediates stress granule-dependent survival during chemotoxic stress. Adv Biol Regul 71:156–171.

Glauninger H, Wong Hickernell CJ, Bard JAM, Drummond DA (2022) Stressful steps: Progress and challenges in understanding stress-induced mRNA condensation and accumulation in stress granules. Mol Cell 82:2544–2556.

Guillén-Boixet J et al. (2020) RNA-Induced Conformational Switching and Clustering of G3BP Drive Stress Granule Assembly by Condensation. Cell 181:346-361.e17

Harding HP, Novoa I, Zhang Y, Zeng H, Wek R, Schapira M, Ron D (2000a) Regulated translation initiation controls stress-induced gene expression in mammalian cells. Mol Cell 6:1099–1108.

Harding HP, Zhang Y, Bertolotti A, Zeng H, Ron D (2000b) Perk Is Essential for Translational Regulation and Cell Survival during the Unfolded Protein Response. Mol Cell 5:897–904.

Hofmann S, Kedersha N, Anderson P, Ivanov P (2021) Molecular mechanisms of stress granule assembly and disassembly. Biochimica et Biophysica Acta (BBA) - Molecular Cell Research 1868:118876

Jain S, Wheeler JR, Walters RW, Agrawal A, Barsic A, Parker R (2016) ATPase-Modulated Stress Granules Contain a Diverse Proteome and Substructure. Cell 164:487–498

Kamelgarn M, Chen J, Kuang L, Jin H, Kasarskis EJ, Zhu H (2018) ALS mutations of FUS suppress protein translation and disrupt the regulation of nonsense-mediated decay. Proceedings of the National Academy of Sciences 115:E11904–E11913

Karran P, Brem R (2016) Protein oxidation, UVA and human DNA repair. DNA Repair (Amst) 44:178–185.

Kedersha N, Chen S, Gilks N, Li W, Miller IJ, Stahl J, Anderson P (2002) Evidence That Ternary Complex (eIF2-GTP-tRNA i Met)-Deficient Preinitiation Complexes Are Core Constituents of Mammalian Stress Granules. Mol Biol Cell 13:195–210.

Kedersha N, Cho MR, Li W, Yacono PW, Chen S, Gilks N, Golan DE, Anderson P (2000) Dynamic shuttling of TIA-1 accompanies the recruitment of mRNA to mammalian stress granules. J Cell Biol 151:1257–1268.

Kedersha N, Panas MD, Achorn CA, Lyons S, Tisdale S, Hickman T, Thomas M, Lieberman J, McInerney GM, Ivanov P (2016) G3BP–Caprin1–USP10 complexes mediate stress granule condensation and associate with 40S subunits. J Cell Biol 212:845–860.

Kedersha N, Stoecklin G, Ayodele M, Yacono P, Lykke-Andersen J, Fritzler MJ, Scheuner D, Kaufman RJ, Golan DE, Anderson P (2005) Stress granules and processing bodies are dynamically linked sites of mRNP remodeling. J Cell Biol 169:871–884.

Kedersha NL, Gupta M, Li W, Miller I, Anderson P (1999) RNA-Binding Proteins Tia-1 and Tiar Link the Phosphorylation of Eif-2α to the Assembly of Mammalian Stress Granules. J Cell Biol 147:1431 LP – 1442.

Khong A, Matheny T, Jain S, Mitchell SF, Wheeler JR, Parker R (2017) The Stress Granule Transcriptome Reveals Principles of mRNA Accumulation in Stress Granules. Mol Cell 68:808-820.e5

Kroschwald S, Maharana S, Mateju D, Malinovska L, Nüske E, Poser I, Richter D, Alberti S (2015) Promiscuous interactions and protein disaggregases determine the material state of stress-inducible RNP granules. Elife 4:e06807.

Lester E, Ooi FK, Bakkar N, Ayers J, Woerman AL, Wheeler J, Bowser R, Carlson GA, Prusiner SB, Parker R (2021) Tau aggregates are RNA-protein assemblies that mislocalize multiple nuclear speckle components. Neuron 109:1675-1691.e9

Li YR, King OD, Shorter J, Gitler AD (2013) Stress granules as crucibles of ALS pathogenesis. Journal of cell biology 201:361–372.

Liu-Yesucevitz L, Bilgutay A, Zhang Y-J, Vanderweyde T, Citro A, Mehta T, Zaarur N, McKee A, Bowser R, Sherman M, Petrucelli L, Wolozin B (2010) Tar DNA binding protein-43 (TDP-43) associates with stress granules: analysis of cultured cells and pathological brain tissue. PLoS One 5:e13250.

Luo Y, Na Z, Slavoff SA (2018) P-Bodies: Composition, Properties, and Functions. Biochemistry 57:2424–2431.

Markmiller S, Soltanieh S, Server KL, Mak R, Jin W, Fang MY, Luo E-C, Krach F, Yang D, Sen A, Fulzele A, Wozniak JM, Gonzalez DJ, Kankel MW, Gao F-B, Bennett EJ, Lécuyer E, Yeo GW (2018) Context-Dependent and Disease-Specific Diversity in Protein Interactions within Stress Granules. Cell 172:590-604.e13.

Marmor-Kollet H et al. (2020) Spatiotemporal Proteomic Analysis of Stress Granule Disassembly Using APEX Reveals Regulation by SUMOylation and Links to ALS Pathogenesis. Mol Cell 80:876-891.e6.

Mateju D, Chao JA (2022) Stress granules: regulators or by-products? FEBS J 289:363–373.

Mateju D, Eichenberger B, Voigt F, Eglinger J, Roth G, Chao JA (2020) Single-Molecule Imaging Reveals Translation of mRNAs Localized to Stress Granules. Cell 183:1801-1812.e13.

Matheny T, van Treeck B, Huynh TN, Parker R (2021) RNA partitioning into stress granules is based on the summation of multiple interactions. RNA 27:174–189

McEwen E, Kedersha N, Song B, Scheuner D, Gilks N, Han A, Chen J-J, Anderson P, Kaufman RJ (2005) Heme-regulated inhibitor kinase-mediated phosphorylation of eukaryotic translation initiation factor 2 inhibits translation, induces stress granule formation, and mediates survival upon arsenite exposure. Journal of Biological Chemistry 280:16925–16933.

Moutaoufik MT, el Fatimy R, Nassour H, Gareau C, Lang J, Tanguay RM, Mazroui R, Khandjian EW (2014) UVC-induced stress granules in mammalian cells. PLoS One 9:e112742–e112742.

Namkoong S, Ho A, Woo YM, Kwak H, Lee JH (2018) Systematic Characterization of Stress-Induced RNA Granulation. Mol Cell 70:175-187.e8.

Nover L, Scharf K-D, Neumann D (1989) Cytoplasmic heat shock granules are formed from precursor particles and are associated with a specific set of mRNAs. Mol Cell Biol 9:1298– 1308.

Padrón A, Iwasaki S, Ingolia NT (2019) Proximity RNA Labeling by APEX-Seq Reveals the Organization of Translation Initiation Complexes and Repressive RNA Granules. Mol Cell 75:875-887.e5.

Park Y-J, Choi DW, Cho SW, Han J, Yang S, Choi CY (2020) Stress granule formation attenuates RACK1-mediated apoptotic cell death induced by morusin. Int J Mol Sci 21:5360.

Pothof J, Verkaik NS, Hoeijmakers JHJ, van Gent DC (2009) MicroRNA responses and stress granule formation modulate the DNA damage response. Cell Cycle 8:3462–3468

Proud CG (2019) Phosphorylation and Signal Transduction Pathways in Translational Control. Cold Spring Harb Perspect Biol 11.

Reineke LC, Cheema SA, Dubrulle J, Neilson JR (2018) Chronic starvation induces noncanonical pro-death stress granules. J Cell Sci 131.

Rhine K, Al-Azzam N, Yu T, Yeo GW (2022) Aging RNA granule dynamics in neurodegeneration. Front Mol Biosci 9.

Riggs CL, Kedersha N, Ivanov P, Anderson P (2020) Mammalian stress granules and P bodies at a glance. J Cell Sci 133.

Roy S (2017) Impact of UV Radiation on Genome Stability and Human Health. In: Ultraviolet Light in Human Health, Diseases and Environment (Ahmad SI, ed), pp 207–219. Cham: Springer International Publishing.

Russo A, Scardigli R, la Regina F, Murray ME, Romano N, Dickson DW, Wolozin B, Cattaneo A, Ceci M (2017) Increased cytoplasmic TDP-43 reduces global protein synthesis by interacting with RACK1 on polyribosomes. Hum Mol Genet 26:1407–1418

Sanders DW, Kedersha N, Lee DSW, Strom AR, Drake V, Riback JA, Bracha D, Eeftens JM, Iwanicki A, Wang A, Wei M-T, Whitney G, Lyons SM, Anderson P, Jacobs WM, Ivanov P, Brangwynne CP (2020) Competing Protein-RNA Interaction Networks Control Multiphase Intracellular Organization. Cell 181:306-324.e28.

Srivastava SP, Kumar KU, Kaufman RJ (1998) Phosphorylation of Eukaryotic Translation Initiation Factor 2 Mediates Apoptosis in Response to Activation of the Double-stranded RNA-dependent Protein Kinase*. Journal of Biological Chemistry 273:2416–2423.

Szaflarski W, Fay MM, Kedersha N, Zabel M, Anderson P, Ivanov P (2016) Vinca alkaloid drugs promote stress-induced translational repression and stress granule formation. Oncotarget 7:30307.

Teixeira D, Sheth U, Valencia-Sanchez MA, Brengues M, Parker R (2005) Processing bodies require RNA for assembly and contain nontranslating mRNAs. Rna 11:371–382.

Tourrière H, Chebli K, Zekri L, Courselaud B, Blanchard JM, Bertrand E, Tazi J (2003) The RasGAP-associated endoribonuclease G3BP assembles stress granules. Journal of Cell Biology 160:823–831

van Treeck B, Protter DSW, Matheny T, Khong A, Link CD, Parker R (2018) RNA self-assembly contributes to stress granule formation and defining the stress granule transcriptome. Proceedings of the National Academy of Sciences 115:2734–2739

Vanderweyde T, Apicco DJ, Youmans-Kidder K, Ash PEA, Cook C, da Rocha EL, Jansen-West K, Frame AA, Citro A, Leszyk JD (2016) Interaction of tau with the RNA-binding protein TIA1 regulates tau pathophysiology and toxicity. Cell Rep 15:1455–1466.

Vanderweyde T, Yu H, Varnum M, Liu-Yesucevitz L, Citro A, Ikezu T, Duff K, Wolozin B (2012) Contrasting pathology of the stress granule proteins TIA-1 and G3BP in tauopathies. Journal of Neuroscience 32:8270–8283.

Vu L, Ghosh A, Tran C, Tebung WA, Sidibé H, Garcia-Mansfield K, David-Dirgo V, Sharma R, Pirrotte P, Bowser R, vande Velde C (2021) Defining the Caprin-1 Interactome in Unstressed and Stressed Conditions. J Proteome Res 20:3165–3178

Wek RC (2018) Role of eIF2α Kinases in Translational Control and Adaptation to Cellular Stress. Cold Spring Harb Perspect Biol 10.

Wek SA, Zhu S, Wek RC (1995) The histidyl-tRNA synthetase-related sequence in the eIF-2 alpha protein kinase GCN2 interacts with tRNA and is required for activation in response to starvation for different amino acids. Mol Cell Biol 15:4497–4506

Wheeler JR, Matheny T, Jain S, Abrisch R, Parker R (2016) Distinct stages in stress granule assembly and disassembly Nilsen TW, ed. Elife 5:e18413

Wolozin B (2014) Physiological protein aggregation run amuck: stress granules and the genesis of neurodegenerative disease. Discov Med 17:47.

Wolozin B, Ivanov P (2019) Stress granules and neurodegeneration. Nat Rev Neurosci 20:649–666.

Yang P, Mathieu C, Kolaitis R-M, Zhang P, Messing J, Yurtsever U, Yang Z, Wu J, Li Y, Pan Q, Yu J, Martin EW, Mittag T, Kim HJ, Taylor JP (2020) G3BP1 Is a Tunable Switch that Triggers Phase Separation to Assemble Stress Granules. Cell 181:325-345.e28.

Ying S, Khaperskyy DA (2020) UV damage induces G3BP1-dependent stress granule formation that is not driven by mTOR inhibition-mediated translation arrest. J Cell Sci 133:jcs248310.

Youn J-Y, Dyakov BJA, Zhang J, Knight JDR, Vernon RM, Forman-Kay JD, Gingras A-C (2019) Properties of Stress Granule and P-Body Proteomes. Mol Cell 76:286–294.

Zhang P, Fan B, Yang P, Temirov J, Messing J, Kim HJ, Taylor JP (2019) Chronic optogenetic induction of stress granules is cytotoxic and reveals the evolution of ALS-FTD pathology. Elife 8:e39578.

