## Supplementary Methods for "The Enigma of UV Stress Granules: Research Challenges and New Perspectives"

### *Supplementary Material*

#### 1 Supplementary Materials and Methods

##### 1.1 Cell Culture and Treatments

U2OS and HaCaT cells were maintained at 37ºC in a CO2 incubator at 5% CO2 in DMEM (Gibco) supplemented with 10% FBS, 1% penicillin/streptomycin, and glutamax (Gibco) (1% for U2OS, 0.2% for HaCaT). For SG induction, cells were treated with sodium arsenite (Sigma Aldrich) or irradiated with UV-C on a Staragene Stratalinker (15 $J/m^2$) for U2OS and 3000 $J/m^2$ for HaCaT). For UV treatment cells were grown to 70-90% confluency in a 12-well plate, media was removed, cells were rinsed with 1X PBS, irradiated with UV and then pre-conditioned media was placed back on the cells. Cells were then prepared for immunofluorescence microscopy.

##### 1.2 Immunofluorescence

1.5 x $10^5$ cells were plated onto glass cover slips in a 12-well plate, and the following day treated as indicated. After SG induction cells were prepared for microscopy as previously described (Fay et al., 2021). Briefly, cells were fixed with 4% paraformaldehyde for 10 minutes, permeabilized with methanol for 10 minutes and then placed in 5% bovine serum albumin (BSA) diluted in 1X PBS to block for at least one hour. Primary antibodies were diluted in blocking solution and placed on the cells to incubate for at least 1 hr at room temperature or overnight at 4ºC. Antibodies against G3BP1 were rabbit-anti-G3BP1 CAT#:13057-2-AP and mouse-anti-G3BP1 CAT #:66486-1-Ig, (Proteintech, both used at 1:2000). Antibodies against RACK1 were rabbit-anti-RACK1 CAT#:27592-1-AP and mouse-anti-RACK1 CAT#: 66940-1-Ig (Proteintech, both used at 1:1000). After washing 3 times in 1X PBS, secondary antibodies were applied and incubated at room temperature for 1 hr: Alexa Fluor® 488 (715-545-150) donkey anti-mouse IgG, Alexa Fluor® 488 (711-545-152) donkey anti-rabbit IgG, Alexa Fluor® 594 (715-585-150) donkey anti-mouse IgG, or Alexa Fluor® 594 (711-585-152) donkey anti-rabbit IgG (Jackson ImmunoResearch, at 1:500 – 1:1000). Coverslips were then mounted using vinol mounting medium (Kedersha et al., 2008) onto microscope slides for imaging.

##### 1.3 Fluorescence *In Situ* Hybridization (FISH)

Cells were plated and treated as described above, then processed for FISH essentially as described (Farny et al., 2009). Briefly, cells were fixed in 4% paraformaldehyde for 10 minutes, then 0.1% Trixon-X100 for 10 minutes, then 70% ethanol for 10 minutes. Cells were equilibrated in 2X saline sodium citrate and then hybridized in hybridization buffer with Cy3-oligo$(dT)_{40}$ probe (IDT DNA) for 1 h at 42°C. Cells were washed three times with 2X SSC for 10 min at room temperature, then processed for immunofluorescences as indicated above (starting from the primary antibody step).

##### 1.4 Image Analysis

Coverslips were observed under the Zeiss AXIO Observer A1 inverted fluorescence microscope at 40X magnification. Cover slips were imaged using a Kiralux 2.3 MP Monochrome CMOS Camera (THORLABS CS235MU), and acquired with MicroManager1.3 (micro-manager.org) open-source software. Image analysis was performed using ImageJ (NIH) as previously described (Aulas et al., 2017).

#### 2 Supplementary References

Farny, N. G., Kedersha, N. L., & Silver, P. A. (2009). Metazoan stress granule assembly is mediated by P-eIF2α-dependent and-independent mechanisms. *Rna*, *15*(10), 1814–1821.

Fay, M. M., Columbo, D., Cotter, C., Friend, C., Henry, S., Hoppe, M., Karabelas, P., Lamy, C., Lawell, M., Monteith, S., Noyes, C., Salerno, P., Wu, J., Zhang, H. M., Anderson, P. J., Kedersha, N., Ivanov, P., & Farny, N. G. (2021). Bisphenol A promotes stress granule assembly and modulates the integrated stress response. *Biology Open*, *10*(1), bio057539. https://doi.org/10.1242/bio.057539

Kedersha, N., Tisdale, S., Hickman, T., & Anderson, P. (2008). Real-time and quantitative imaging of mammalian stress granules and processing bodies. *Methods in Enzymology*, *448*, 521–552.
